# Self-powered electronics-free Wearable Disposable Electrotherapy (WDE) platform for accelerated wound healing

**DOI:** 10.64898/2026.07.22.740124

**Authors:** Mojtaba Belali Koochesfahani, Kyle Donnery, Abhiramalakshmi Senthil Kumar, Rayyan Bhuiyan, Derek A. Drumm, Baseer Mirza, Miguel Posada Perez, Miguel R. Diaz Uraga, Guillermo Núñez Ponasso, Sergey N. Makaroff, Mohamad FallahRad, Marom Bikson

## Abstract

Electrical stimulation accelerates wound repair by modulating endogenous bioelectric signals that regulate inflammation, angiogenesis, extracellular matrix remodeling, and cellular responses within the wound microenvironment. However, clinical translation has been hindered by cumbersome devices with procedures that disrupt standard wound-care workflows, direct electrode contact with the wound bed, and/or limited stimulation output. Wearable Disposable Electrotherapy (WDE) integrates an electronics-free printed electrochemical architecture into an mm-thick patch that looks and is applied like a conventional bandage. The device is self-powered and delivers a single electrotherapy dose simply by application to the skin. Device dose-control (electrochemical performance) and efficacy were evaluated in a full-thickness excisional wound model in rats, compared with a sham device and a conventional Constant Current (CC) stimulator. WDE or control treatments were applied daily from day 1 through day 13, with endpoint evaluation on day 14. WDE delivered electrical stimulation comparable to CC while reducing the time required to achieve 50% wound closure by 2.08 days (∼29%) relative to sham treatment. Histological and immunofluorescence analyses at day 14 demonstrated enhanced tissue remodeling, including increased collagen deposition (∼25%), tissue cellularity (∼73%), myofibroblast-associated αSMA expression (∼2.6-fold), angiogenesis-associated CD31 expression (∼2.0-fold), and increased expression of both M2- (CD206, ∼2.0-fold) and M1- associated (iNOS, ∼1.7-fold) markers compared with sham. A novel cellular-resolution dosimetry model, leveraging charge-based boundary element method accelerated with the fast multipole method (BEM-FMM), provides a biophysical framework linking electrical stimulation with wound microenvironment and tissue repair mechanisms. Together, these findings establish WDE as a practical bioelectric wound dressing that accelerates wound healing and tissue remodeling, with the simplicity and scalability of disposable bandages.

## Introduction

Wound healing depends on the orchestration of inflammation, angiogenesis, extracellular matrix remodeling, and tissue regeneration [1,2]. Alongside biochemical signaling, endogenous electric fields regulate these regenerative processes [3–5]. Ion transport across intact epithelium establishes a transepithelial potential that is redistributed following injury, generating lateral electric fields at the wound margins [3]. These endogenous bioelectric cues direct the migration (electrotaxis), proliferation, and differentiation of keratinocytes, fibroblasts, endothelial cells, and immune cells [4–7]. Exogenous electrical stimulation reinforces these physiological signals, accelerating wound closure and promoting angiogenesis, collagen deposition, pro-repair inflammatory responses, and improved tissue regeneration across experimental and clinical studies [7–9].

Despite broad evidence for wound electrotherapy, clinical adoption is limited by usability [7,10]. Existing devices require wiring the dressing to an external stimulator or are self-contained but provide low (∼10 µA) and/or passively governed (non-programmable) stimulation output [9–12]. Moreover, these devices mechanically disrupt the wound bed or clinical wound-care workflows [10,11,13]. These usability factors impede adoption across routine (e.g., home care) and medically involved (e.g., postoperative) care.

To address translational barriers across electrotherapy, we developed the Wearable Disposable Electrotherapy (WDE) platform: an electronics-free (environmentally benign), self-powered single-programmed-dose electrotherapy technology based on a printed 3D electrochemical architecture integrated into disposable mm-thick adhesive patches [11]. Here, we adapt WDE for wound healing by redesigning the electrode configuration to deliver lateral direct-current stimulation across the wound through electrodes on intact peri-wound skin (thereby avoiding direct electrode contact with the wound bed and repeated mechanical disruption), scaling the electrical dose for cutaneous applications, and implementing interchangeable device geometries suitable for a wide range of wound sizes. The resulting devices retain the form-factor and application of conventional disposable bandages while delivering a defined single-use electrotherapy dose.

In this study, we engineered the WDE platform for wound electrotherapy and evaluated its electrical performance, therapeutic efficacy, and mechanisms of action in a full-thickness rat wound model. Performance was benchmarked against a sham device and conventional constant-current stimulation. Wound repair was quantified through longitudinal healing, histological and immunofluorescence analyses of tissue remodeling and regenerative responses, including angiogenesis and inflammatory responses, together with electrochemical characterization and cellular-resolution computational modeling of stimulation biophysics within the wound microenvironment. These studies evaluate whether a self-powered, bandage-format electrotherapy platform can combine therapeutic electrical stimulation with a clinically practical form factor.

## Methods

### Device design, fabrication, and electrochemical characterization

We engineered a wound electrotherapy device leveraging our Wearable Disposable Electrotherapy (WDE) multilayer printed technology [11]. The flexible self-powered device integrates a printed electrochemical battery, stimulation electrodes, conductive hydrogel interfaces, adhesive layers, and wound dressing into a disposable electrotherapy platform. Device geometries were adapted to accommodate different wound sizes while preserving the same peri-wound electrode configuration and electrical operating principle. The WDE fabrication process and constituent materials are scalable and environmentally benign [11], with the device internal architecture and overall geometries and electrical dose designed for wound healing.

Electrochemical performance of the stimulation electrodes was evaluated under galvanostatic discharge conditions using an ionic bridge consisting of foam saturated with isotonic saline (0.9% NaCl). Hydrogel electrodes were placed in contact with foam, representing a hydrated ionic interface encountered during device operation. Individual Ag/AgCl cathodes and zinc anodes were connected to a PalmSens4 (PalmSens BV, Houten, The Netherlands) operated in galvanostatic mode and discharged at a constant current of 300 µA for 120 minutes, while electrode potentials were continuously recorded [11]. Electrochemical characterization demonstrated stable galvanostatic performance of the stimulation electrodes over the intended 120-minute treatment period (Supplementary Figure 1A). Under 300 µA constant-current discharge, the Ag/AgCl stimulation electrode exhibited minimal polarization, whereas the zinc stimulation electrode generated progressive anodic polarization with a gradual decrease in potential, consistent with progressive oxidation during galvanostatic discharge.

Electrode-skin impedance characterization during dose delivery was performed using the hydrogel stimulation electrode pair. Hydrogel electrodes were positioned on intact skin immediately rostral and caudal to the wound, avoiding direct mechanical contact between the stimulation electrodes and the wound bed. Stimulation sessions were performed while animals were awake and freely moving within their cages. Electrical stimulation was delivered using a custom analog front-end interfaced with an Analog Discovery 3 multifunction instrument (Digilent Inc., Pullman, WA, USA), configured to deliver a constant current of 300 µA throughout 120 minutes. Output voltage was continuously recorded (Supplementary Figure 1B).

The printed battery was independently characterized by galvanostatic discharge at 300 µA for 120 minutes using a Keithley 2450 SourceMeter SMU (Keithley Instruments, Cleveland, OH, USA), with battery voltage continuously recorded throughout the test. Independent characterization of the printed battery confirmed sustained output throughout the targeted treatment duration (Supplementary Figure 1C). During a 300 µA constant-current discharge, battery voltage gradually decreased from ∼1.6 to ∼1.1 V over 120 minutes, maintaining sufficient potential to support continuous electrotherapy.

For in vivo electrical characterization, voltage and current delivered by both the WDE and external constant-current (CC) systems were continuously recorded during each 120-minute stimulation session using a custom analog front-end interfaced with an Analog Discovery 3 multifunction measurement instrument (Digilent Inc., Pullman, WA, USA). The delivered charge was calculated by numerical integration of the measured current over time.

The surface morphology of the printed battery and stimulation electrodes was characterized using scanning electron microscopy (SEM; SUPRA 55, Carl Zeiss, Oberkochen, Germany) operated at an accelerating voltage of 16 kV using a backscattered electron detector. Samples were imaged without conductive coating. Scanning electron microscopy (SEM) revealed characteristic surface morphologies of the printed electrochemical (battery and stimulation electrode) components (Supplementary Figure 1D). The printed zinc anode exhibited a particulate morphology, whereas the cathode showed a denser aggregated structure. The Ag/AgCl stimulation electrode displayed a relatively uniform surface, while the zinc stimulation electrode exhibited the rougher morphology characteristic of electroplated zinc.

The physiological load impedance was measured under constant-current (CC) stimulation for the intended dose (300 µA, 120-minute treatment). Electrode-skin impedance (calculated from the measured voltage) gradually decreased from ∼6.3 kΩ at the onset of stimulation to ∼3.3 kΩ at 90 minutes, consistent with progressive hydrogel hydration and improved electrical coupling at the tissue-electrode interface [14].

The fabricated WDE for rodent-model wound electrotherapy integrates the optimized printed battery, stimulation electrodes, conductive hydrogel interfaces, adhesive layers, and wound dressing into a thin, single-dose, flexible disposable patch (Supplementary Figure 1E). Electrical performance of the complete fabricated WDE platform for rodent wound electrotherapy was evaluated during in vivo stimulation (Supplementary Figure 1F). The device delivered an average current of 286 ± 98 µA, not significantly different from the 300 µA target - as would be provided by a CC stimulator (Welch’s *t*-test, *p* > 0.05). Delivered charge was likewise comparable (CC: 2.16 C; WDE: 2.06 ± 0.71 C; *p* > 0.05). WDE current output remained stable throughout the treatment period before gradually declining near completion, consistent with the designed progressive depletion of the printed battery and increased internal resistance [11].

### Experimental wound healing model

All experimental procedures were approved by the Institutional Animal Care and Use Committee (IACUC) of The City College of New York (Protocol No. 2024-0003) and were conducted in accordance with the NIH Guide for the Care and Use of Laboratory Animals. A full-thickness excisional wound model was used based on established preclinical approaches for studying cutaneous wound healing [15]. Male Sprague-Dawley rats (7-9 weeks old, 250-300 g; Charles River Laboratories, Wilmington, MA, USA) were used for all experiments. Animals were housed under standard laboratory conditions with ad libitum access to food and water and allowed to acclimate for up to 4 days before experimentation.

On day 1, following dorsal skin preparation under general anesthesia, a full-thickness excisional wound (10 mm diameter or two 6 mm diameter) was excised on the dorsal midline using a sterile biopsy punch. Baseline wound images were acquired immediately after surgery. Animals were then randomly assigned to one of three experimental groups: Wearable Disposable Electrotherapy (WDE; n = 13), constant-current stimulation (CC; n = 12), or sham (n = 13) device treatment. No animals were excluded from the longitudinal wound-healing analysis.

Electrical stimulation and wound imaging were performed once daily from day 1 through day 13, according to the assigned treatment protocol. At the conclusion of the study (day 14), final imaging was done, animals were euthanized, and wound tissues were collected for histological and immunofluorescence analyses.

### Experimental design and stimulation protocols

In all groups, stimulation or sham was delivered using a peri-wound electrode configuration in which conductive hydrogel electrodes were positioned on intact skin immediately rostral and caudal to the wound, avoiding direct mechanical contact between the stimulation electrodes and the wound bed.

Electrical stimulation was applied once daily for 120 minutes from day 1 through day 13, followed by wound imaging and tissue collection on day 14. Stimulation sessions were performed while animals were awake and freely moving within their cages. In the sham group, the device was applied using the identical electrode configuration without active electrical stimulation.

In the CC group, electrical stimulation was delivered using a custom analog front-end interfaced with an Analog Discovery 3 multifunction instrument (Digilent Inc., Pullman, WA, USA), configured for a constant current of 300 µA, corresponding to the nominal target current of the WDE device. In the WDE group, stimulation was delivered using the self-powered wearable disposable electrotherapy device, which generated an average current of 286 ± 98 µA through the 120-minute stimulation period.

Following each stimulation session, the stimulation components were removed while the wound dressing remained in place to protect the wound between treatments. Fresh conductive hydrogel electrodes and WDE devices were applied before each subsequent daily stimulation session.

Electrical output (voltage and current) was continuously monitored during all stimulation sessions. The delivered charge was calculated by integrating the measured current over time, and electrode-skin impedance was estimated from the measured voltage and current using Ohm’s law.

### Wound imaging and quantification

Wound healing progression was monitored by daily digital imaging. Images were acquired immediately following wound creation (day 1) and once daily thereafter using a digital camera (MU300, AmScope, Irvine, CA, USA) mounted on a stereomicroscope (SZ61, Olympus Corporation, Tokyo, Japan). Image acquisition was performed using AmScope software (version 4.12, AmScope).

Longitudinal wound area was quantified using a semi-automated image segmentation pipeline with manual validation. A subset of wound images was manually annotated using a custom graphical user interface (GUI) developed in MATLAB (MathWorks, Natick, MA, USA; version R2025b), in which wound boundaries were delineated by visual inspection. These manually generated masks served as reference annotations for automated wound segmentation across the complete image dataset. The wound area was calculated from the resulting segmentation masks and normalized using a 2-mm circular calibration marker included in each image.

To evaluate segmentation accuracy and minimize observer bias, an independent manual analysis was then performed. Two blinded investigators independently delineated wound boundaries using QuPath (version 0.6.0), and wound areas were calculated from the resulting annotations using custom Python scripts (Python version 3.14). Measurements obtained from the semi-automated and manual analyses were compared to verify the robustness of the segmentation pipeline.

Wound area at each time point was normalized to the corresponding wound area immediately after surgery (day 1) and expressed as relative wound size (%) according to Equation 1.

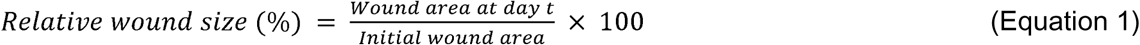

Visible wound exudate was assessed from daily wound dressing changes by visual inspection and recorded as present or absent for each animal at each time point. Time to exudate resolution was defined as the first day on which visible exudate was absent and remained absent thereafter. Wounds with persistent visible exudate at the final day-14 observation were assigned a value of day 15 for calculation of the estimated exudate-resolution time.

### Histological analysis

At day 14, full-thickness wound tissues were excised using a sterile biopsy punch and rinsed in phosphate-buffered saline (PBS) to remove residual blood. Samples were fixed in 4% paraformaldehyde at 4°C overnight. Following fixation, tissues were washed in PBS (3 × 5 minutes) and cryoprotected in 30% sucrose at 4°C overnight.

Cryoprotected tissues were embedded in optimal cutting temperature (OCT) compound, with the epidermal surface oriented parallel to the sectioning plane. Embedded samples were rapidly frozen and stored at -80°C until cryosectioning.

Cryosections (6 µm) were prepared using a cryostat (CM1950, Leica Microsystems, Wetzlar, Germany) and mounted onto glass microscope slides. Sections were air-dried at room temperature before staining.

General tissue morphology was evaluated using hematoxylin and eosin (H&E) staining performed with a commercial staining kit (ab245880, Abcam, Cambridge, UK). Collagen deposition and connective tissue organization were evaluated using Masson’s Trichrome staining performed with a commercial connective tissue staining kit (ab150686, Abcam) according to the manufacturer’s protocols.

### Immunofluorescence staining

Tissue sections were washed three times in 1× phosphate-buffered saline (PBS) for 10 minutes each and incubated for 1 h at room temperature in blocking buffer consisting of 1× PBS supplemented with 5% donkey serum and 0.2% Triton X-100. Primary antibodies were diluted in 1× PBS and incubated with the tissue sections for 2 h at room temperature or overnight at 4°C in a humidified, light-protected chamber. Sections were subsequently washed three times in 1× PBS for 10 minutes each and incubated with the appropriate fluorescent secondary antibodies for 2 h at room temperature. Following secondary antibody incubation, sections were washed in PBS, counterstained with DAPI, mounted using a fluorescence-compatible mounting medium, and coverslipped. Fluorescent images were acquired using a BZ-X810 fluorescence microscope (Keyence Corporation, Osaka, Japan) under identical imaging settings for all specimens within each staining group.

Primary antibodies included CD31 (BS0468R, Thermo Fisher Scientific, Waltham, MA, USA), α- smooth muscle actin (αSMA; PA5-18292, Thermo Fisher Scientific), inducible nitric oxide synthase (iNOS; PA1-036, Thermo Fisher Scientific), and CD206 (PA5-101657, Thermo Fisher Scientific).

### Histological and immunofluorescence image quantification

Quantitative analysis of histological and immunofluorescence images was performed using QuPath (version 0.6.0) and Fiji (ImageJ version 1.54s). Regions of interest (ROIs) encompassing the wound area and adjacent intact skin were manually defined based on histological landmarks. Quantitative measurements were normalized to the corresponding intact skin region from the same tissue section to minimize inter-sample variability. All quantitative image analyses were performed by investigators blinded to treatment allocation.

Masson’s Trichrome-stained sections were analyzed to quantify relative collagen deposition, whereas hematoxylin and eosin (H&E)-stained sections were used to quantify eosin intensity, epidermal thickness, epidermal junction density, and relative cell density. Fractal dimension and lacunarity were quantified from binarized histological images using ImageJ with the FracLac plugin, following a previously described FracLac-based image-analysis workflow [16]. Identical image-processing and analysis parameters were applied to all specimens.

Immunofluorescence images were analyzed by measuring the mean fluorescence intensity within the wound region for each marker (αSMA, CD31, CD206, and iNOS). Marker expression was expressed as the ratio of fluorescence intensity within the wound region to that of the corresponding intact skin region. The macrophage polarization index was estimated as the ratio of iNOS to CD206 fluorescence intensity (iNOS/CD206).

All image analyses were performed using identical processing parameters for all samples within each experimental group.

### Computational modeling

A two-dimensional computational model was developed to investigate current flow and electric field distributions within the wound microenvironment during electrical stimulation. The model incorporated anatomically representative skin microarchitecture together with heterogeneous tissue electrical properties to evaluate the influence of tissue composition on localized electrical stimulation.

The computational geometry was constructed from representative histological architecture derived from hematoxylin and eosin (H&E)-stained skin sections reported in the literature [17–21]. Anatomically resolved tissue structures were generated from segmented DXF geometries and included the epidermis, dermis, sweat glands and ducts, sebaceous glands, hair follicles, vasculature, nerves, hypodermal adipose tissue, and muscle. Electrical conductivity values were assigned to individual tissue and cellular compartments based on previously reported measurements of dielectric and bioelectrical properties. The principal tissue conductivities were 10^-5^ S/m for the stratum corneum, 0.0091 S/m for the viable epidermis, 0.526 S/m for the dermis, 0.026 S/m for adipose tissue, 0.585 S/m for muscle, 0.662 S/m for blood-filled vessel lumens, 1.0 S/m for sweat ducts, and 0.39 S/m for neural tissue. Additional tissue structures were assigned conductivities of 0.5 S/m for sweat gland regions and blood vessel walls, 0.026 S/m for sebaceous gland regions, and 0.4 S/m for hair walls. Cellular membranes were represented as highly resistive regions with a conductivity of 10^-6^ S/m, whereas cellular nuclei were assigned a conductivity of 0.3 S/m, except for adipocyte nuclei, which were assigned 0.1 S/m. Tissue compartments for which direct experimental conductivity measurements were unavailable were assigned values based on the closest physiological or electrophysiological analogs [22–27].

Electrical current propagation was simulated using a charge-based boundary element method accelerated with the fast multipole method (BEM-FMM), implemented in custom MATLAB scripts (MathWorks, Natick, MA, USA) following previously described bioelectromagnetic modeling frameworks [28,29]. The governing boundary integral equations were solved using a Galerkin formulation with an iterative flexible generalized minimal residual (FGMRES) solver [29,30]. Current was prescribed at boundary regions corresponding to the hydrogel electrode interfaces, with ±0.3 mA distributed uniformly along each electrode boundary by normalizing the applied current to the total electrode-boundary length in the two-dimensional formulation. The resulting boundary charge solution was used to compute current density and electric field distributions throughout the modeled tissue domain. Simulated current density and electric field distributions were subsequently analyzed throughout the wound microenvironment to evaluate the influence of heterogeneous tissue electrical properties on localized current pathways and electrical field distributions surrounding the wound.

### Statistical analysis

Statistical analyses were performed using Python (version 3.14) and MATLAB R2025b (MathWorks, Natick, MA, USA). Primary endpoints were longitudinal wound closure and time to 50% wound closure; histological and immunofluorescence endpoints were exploratory.

Longitudinal wound closure data were analyzed using linear mixed-effects (LME) models implemented with the statsmodels package, with treatment group, time, a quadratic time term (time^2^), and their interactions with treatment included as fixed effects, and animal included as a random intercept to account for repeated measurements. Model selection was based on the Akaike information criterion (AIC), which favored the model including the quadratic time term. Time to 50% wound closure was compared using Welch’s two-sample t-test; across the three pairwise contrasts, Holm-adjusted p-values were 0.003 (WDE vs. sham), 0.004 (CC vs. sham), and 0.24 (CC vs. WDE), which did not alter any conclusion. Pairwise day-by-day comparisons of relative wound size are descriptive and uncorrected; longitudinal inference is based on the LME model.

Histological and immunofluorescence measures are ratio-scale (wound tissue normalized to adjacent intact skin) and were compared between WDE and sham using Welch’s two-sample t-test (n = 4 animals per group). Welch’s formulation was pre-specified because equal variances could not be assumed. A parametric approach was pre-specified; p-values for these endpoints are nominal and uncorrected for multiple comparisons; they describe the magnitude and direction of treatment-associated differences.

Data are presented as mean ± standard deviation (SD) unless otherwise indicated. All statistical tests were two-sided, and differences were considered statistically significant at p < 0.05 for primary endpoints. Comparisons between CC and WDE were not powered as equivalence tests; the absence of a detected difference does not establish equivalence.

A sensitivity analysis was performed for the three-group experimental design using a one-way analysis of variance framework. Using a conservative balanced approximation of 12 animals per group (36 animals total) and a two-sided significance level of α = 0.05, the study provided 80% power to detect an omnibus effect size of Cohen’s f = 0.54, corresponding to a large treatment effect. Because longitudinal wound measurements were analyzed using a linear mixed-effects model with repeated observations within animals, this represents a conservative approximation of the statistical sensitivity of the study. For histological and immunofluorescence endpoints, four animals per group were analyzed; a two-sided independent-samples framework with α = 0.05 indicated 80% power to detect a standardized effect of Cohen’s d = 2.38.

## Results

### Wearable Disposable Electrotherapy (WDE) for wound healing platform

To develop a practical and effective electrotherapy platform for wound care, we adapted the Wearable Disposable Electrotherapy (WDE) platform [11] to deliver peri-wound stimulation using a thin, flexible, self-powered patch (Figure 1A). The multilayer architecture integrates a printed battery, stimulation electrodes, conductive hydrogel interfaces, adhesive layers, and a wound dressing into a compact disposable device. WDE design is based on scalable manufacture, with each device a sealed mm-thick conformable patch containing all power, control, and interface elements (Figure 1 B). The modular architecture supports tailored geometries to accommodate wounds of different sizes while preserving the same peri-wound stimulation strategy (Figure 1C). Hydrogel electrodes positioned on intact peri-wound skin deliver current across the wound while avoiding direct electrode contact with the wound bed. Figure 1D illustrates the generation of tangential current flow by WDE devices across the wound perimeter, microscopic structures that influence electric fields (through tissue heterogeneity), and associated cellular responses enhancing tissue repair.

**Figure 1.**
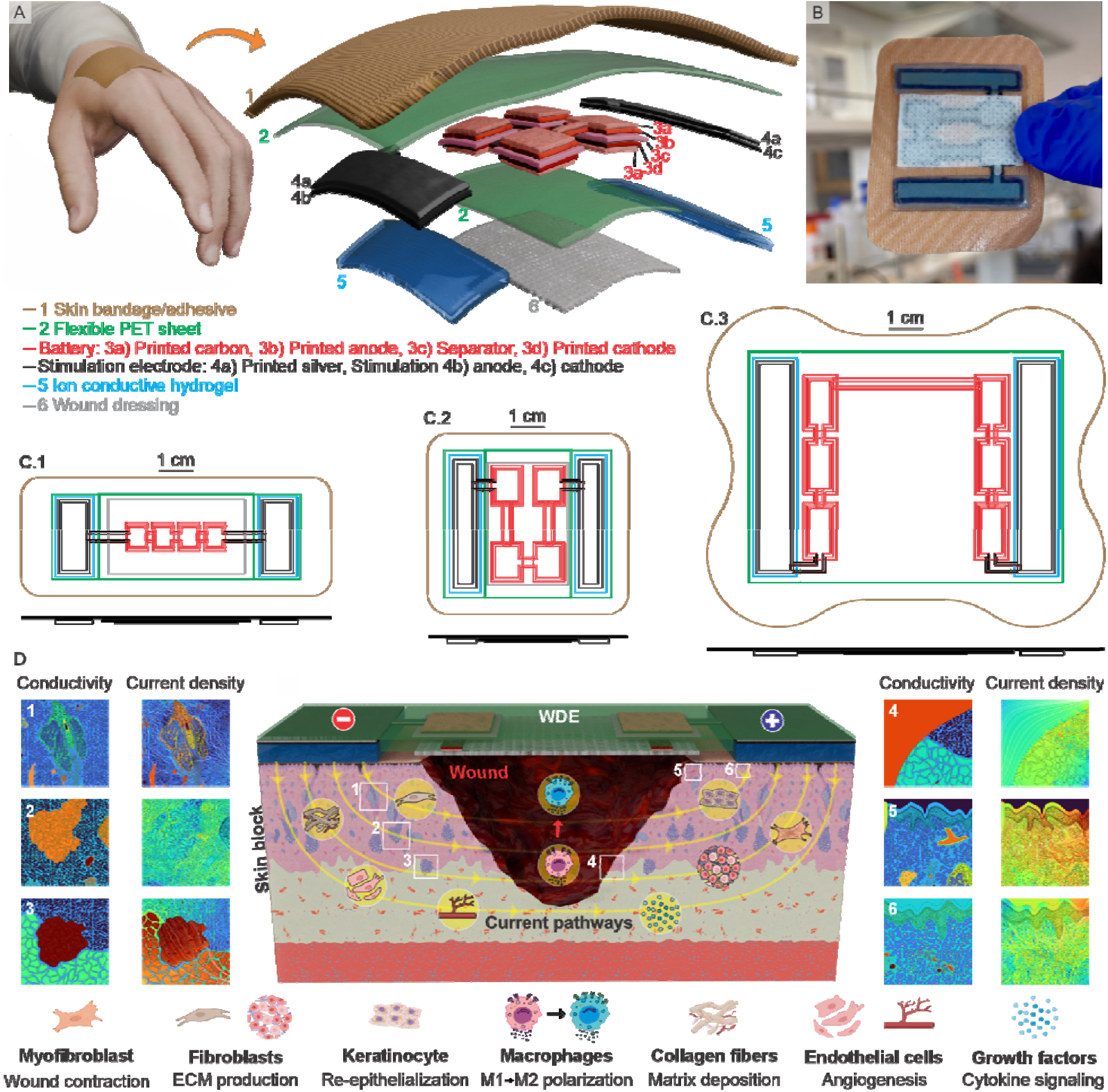
| Design concept, scalable architecture, and proposed mechanism of action of the Wearable Disposable Electrotherapy (WDE) wound healing platform. **(A)** WDE schematic with multilayer architecture. The mm-thick flexible patch is applied like a conventional adhesive dressing. The wound is under the felt, avoiding hydrogel contact and mechanical disruption. Lateral stimulation is delivered between hydrogel electrodes on intact peri-wound skin, which also provides device adhesion. The self-powered battery architecture delivers a defined single-use dose upon application. **(B)** Photograph of the fabricated WDE device (medium size). The thin, flexible form factor conforms to the skin. **(C)** Scalable WDE layouts for small (C.1), medium (C.2), and large (C.3) wound geometries. Modular battery design enables a range of electrical doses applied across various wounds. **(D)** Proposed mechanism of WDE- mediated wound healing illustrating microscopic tissue electrical properties, simulated current path, and representative regenerative cellular responses.

### Design and validation of the Wearable Disposable Electrotherapy device- experimental wound healing model

To evaluate the therapeutic efficacy of the WDE platform, a full-thickness excisional wound healing model was established in rats and compared with constant-current (CC) stimulation and sham treatment (Figure 2). Animals were acclimated for three days before surgery. On day 1 (surgery day), full-thickness excisional wounds were created on the dorsal skin, followed by baseline wound imaging and immediate application of the assigned treatment. Electrotherapy and wound imaging were repeated daily through day 13, with endpoint imaging at day 14, and wound tissues were collected for histological and immunofluorescence analyses (Figure 2A.1). Throughout each stimulation session, voltage and current were continuously monitored to characterize electrical dose delivery (Figure 2A.2).

**Figure 2.**
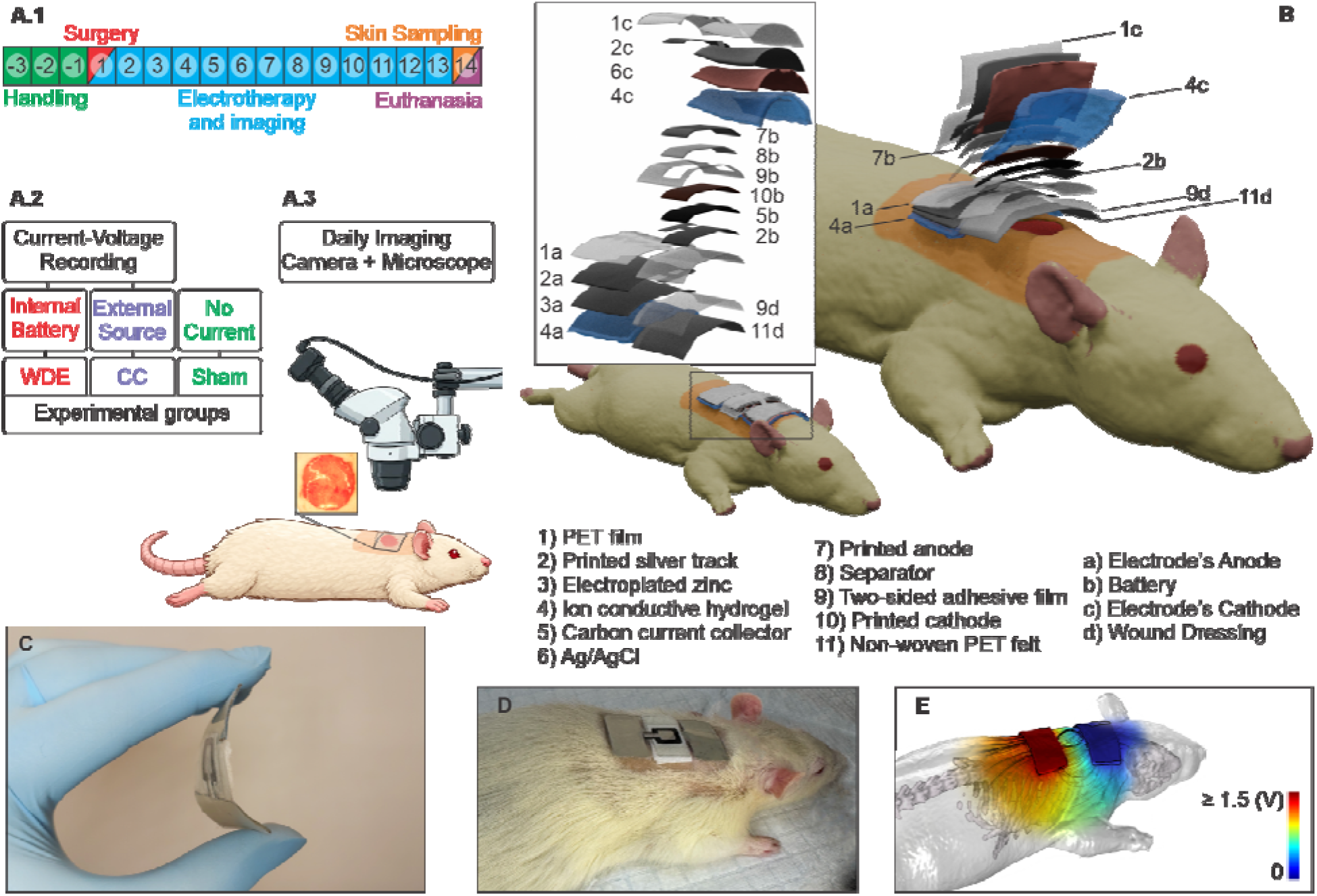
| Wearable Disposable Electrotherapy configured for the experimental wound healing rat model. **(A.1)** Experimental timeline. Rats were acclimated for three days before surgery (days -3 to -1). Full-thickness excisional wounds were created on day 1, followed by daily wound imaging and electrotherapy. Electrotherapy continued daily through day 13. On day 14, final imaging was done, and tissue samples were collected for histological and immunofluorescence analysis. **(A.2)** Schematic of the experimental setup with electrotherapy delivery and recording of stimulation voltage/current during electrotherapy. **(B)** Exploded view of WDE structure. **C)** Photograph of a fabricated device, conforming to the body with a thin profile. **D)** WDE was applied to the dorsal wound of a rat, with peri-wound electrode placement and bandage-format application. **E)** Macroscopic (tissue-level) computational model of the adapted WDE configuration: simulated electric field magnitude (false color) and tangential current density (streamlines) during wound stimulation.

The WDE device was adapted for rodent peri-wound stimulation as a compact multilayer patch integrating the printed battery, stimulation electrodes, conductive hydrogel interfaces, adhesive layers, and wound dressing (Figure 2B-D, See Methods, Supplementary Figure 1). The WDE group received self-powered electrical stimulation, whereas the CC group received a comparable stimulation dose from an external current source, and the sham group received the identical device configuration without electrical stimulation. In all groups, hydrogel electrodes were positioned on intact skin adjacent to the wound, allowing current to pass across the wound bed while avoiding direct electrode contact with injured tissue.

Macroscopic computational modeling of the complete device configuration predicted electric field distribution and current propagation through the wound region during stimulation (Figure 2E). The peri-wound electrode configuration directed current through the wound microenvironment while minimizing electrochemical interactions at the wound surface, reducing localized current hot spots, and preserving the integrity of the healing tissue during repeated device application and removal.

### WDE accelerates wound closure

Representative wound images demonstrated progressive healing in each group over the 14-day study, with faster wound closure in both electrically stimulated groups than in the sham group (Figure 3A.1). Wound boundary overlays further illustrate the accelerated reduction in wound area following both constant-current (CC) stimulation and WDE treatment (Figure 3A.2).

**Figure 3.**
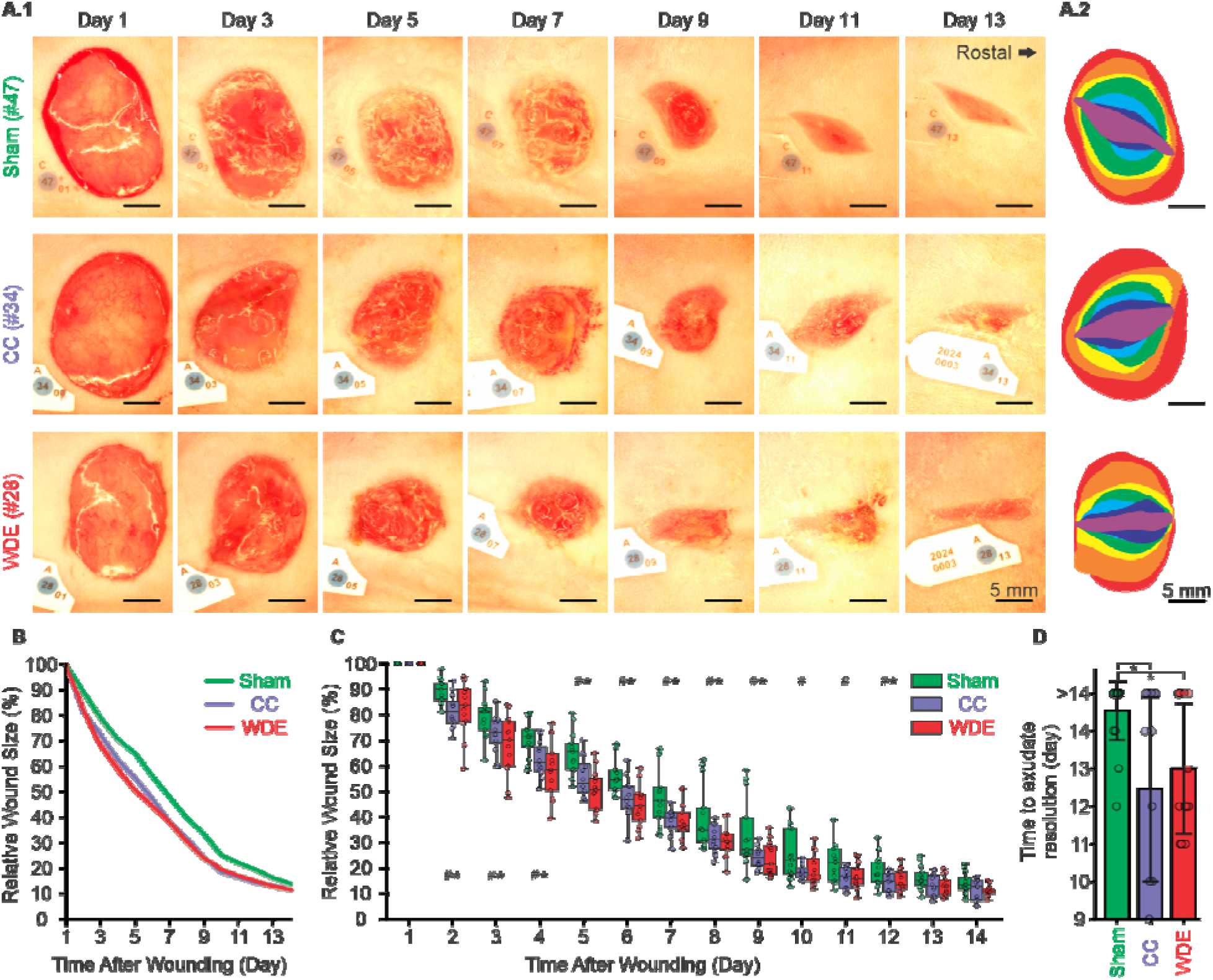
| Accelerated full-thickness wound healing following Wearable Disposable Electrotherapy (WDE). **(A.1)** Representative wound images acquired at days 1, 3, 5, 7, 9, 11, and 13 post-injury from rats treated with a sham device, constant-current (CC) stimulation, or Wearable Disposable Electrotherapy (WDE). Wound areas were quantified referencing a 2-mm circular reference marker. Scale bars, 5 mm. **(A.2)** Corresponding wound boundary overlays. Colors indicate wound margins at successive time points. **(B)** Time course of relative wound size (normalized to day 1) following injury. Solid lines represent group means (shaded regions indicate the range [minimum-maximum] across animals; sham n = 13, WDE n = 13, CC n = 12). A linear mixed-effects model revealed a significant group × time interaction, indicating accelerated wound healing in the CC and WDE groups compared with sham (group × day: p < 0.001; group × day^2^: p < 0.001), whereas no significant difference was observed between CC and WDE groups. **(C)** Distribution of relative wound size across groups for each experimental day (days 1-14). Individual animals are shown with box-and-whisker plots. Pairwise comparisons between groups at each time point were performed using Welch’s t-tests (*WDE vs. sham; #CC vs. sham; p < 0.05). **(D)** Estimated day of complete exudate resolution across treatment groups. (*p < 0.05; bars: mean ± SD; individual animals).

Longitudinal analysis demonstrated significantly accelerated wound closure in both electrically stimulated groups compared with sham (Figure 3B, C). A linear mixed-effects model identified significant treatment × time (p < 0.001) and treatment × time^2^ (p < 0.001) interactions, consistent with faster healing trajectories in the CC and WDE groups when compared to sham. No significant differences were observed between CC and WDE (treatment × time: p = 0.366; treatment × time^2^: p = 0.436), indicating that the self-powered WDE accelerates wound healing comparable to conventional CC stimulation.

The time required to reach 50% wound closure was significantly reduced by electrical stimulation. Sham-treated wounds reached 50% wound closure after 7.10 ± 1.36 days, whereas CC- and WDE-treated wounds reached this milestone after 5.54 ± 0.76 days and 5.02 ± 1.35 days, corresponding to 22% and 29% reductions, respectively (Welch’s t-test with Holm adjustment; CC vs. sham: p = 0.004; WDE vs. sham: p = 0.003). No statistically significant difference was detected between CC and WDE (adjusted p = 0.24).

Healing progression was further assessed by the resolution of visible wound exudate (Figure 3D). Estimated times to complete exudate resolution were 13.46 ± 0.97 days for sham, 11.42 ± 2.47 days for CC, and 12.00 ± 1.73 days for WDE. Relative to sham, exudate resolved approximately 15% earlier with CC and 11% earlier with WDE (Welch’s t-test; CC vs. sham: p = 0.018; WDE vs. sham: p = 0.016), with no significant difference between the two electrically stimulated groups (p = 0.50). By day 14, complete exudate resolution was observed in only 30.8% of sham-treated wounds, compared with 71.4% and 54.5% of CC- and WDE- treated wounds, respectively.

Collectively, these results demonstrate that daily peri-wound electrical stimulation accelerates wound healing by promoting faster wound closure and earlier exudate resolution. The self-powered WDE platform produced therapeutic outcomes comparable to those achieved with a conventional CC stimulator, supporting its potential as a practical disposable electrotherapy platform for wound care.

### WDE enhances tissue remodeling and structural regeneration

Histological evaluation at day 14 showed differences in tissue architecture between WDE- treated and sham wounds (Figure 4). Representative Masson’s Trichrome (MT) and hematoxylin and eosin (H&E) sections revealed more mature tissue organization and improved wound remodeling following WDE treatment (Figure 4A, B).

**Figure 4.**
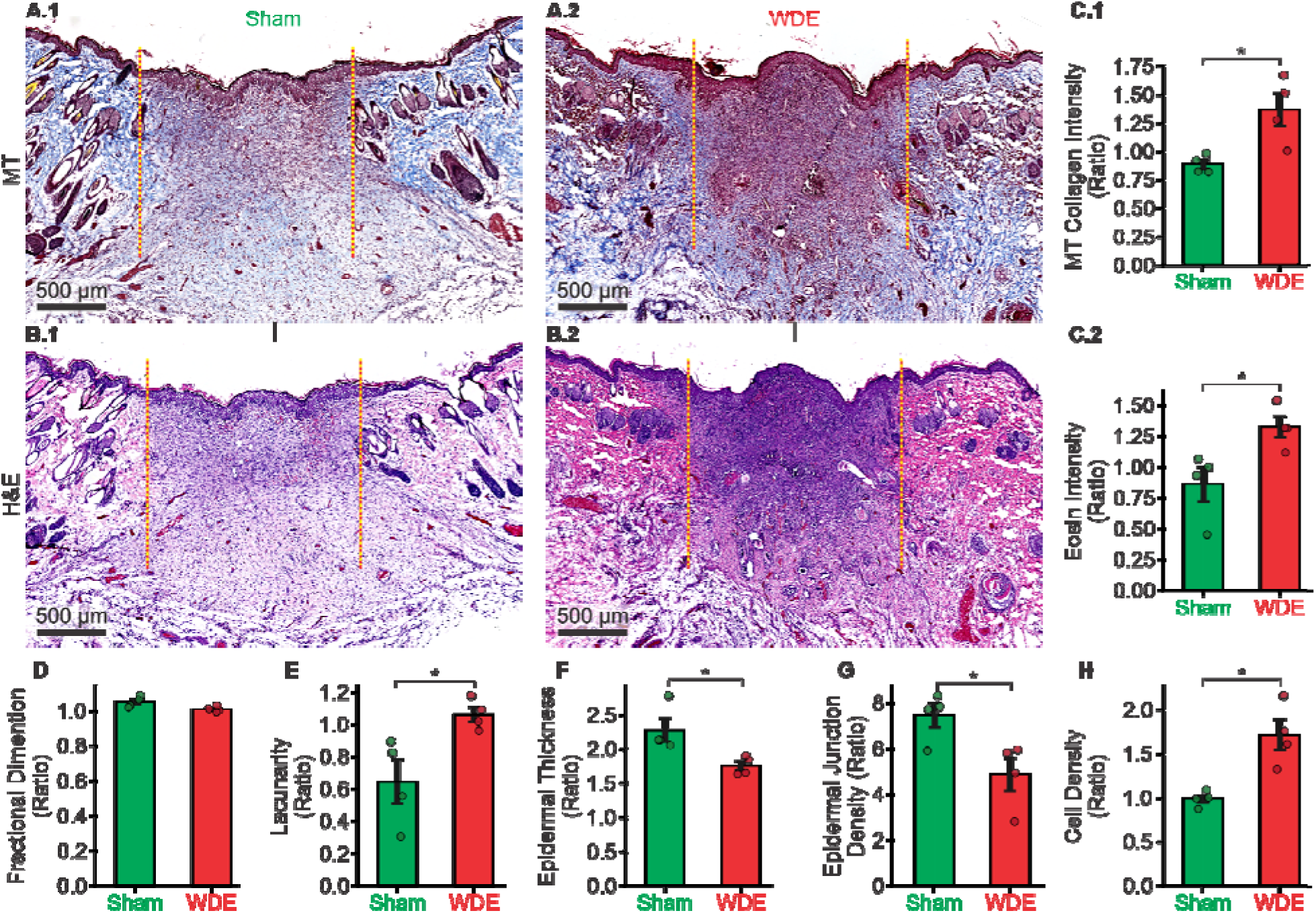
| Histological evaluation of tissue remodeling after 14 days of electrotherapy. **(A.1-A.2)** Representative Masson’s Trichrome (MT)-stained sections from sham- and WDE- treated wounds. **(B.1-B.2)** Corresponding hematoxylin and eosin (H&E)-stained sections. Dashed lines indicate the approximate wound margins. **(C.1-C.2)** Quantification of collagen deposition (MT intensity) and eosin intensity. **(D-H)** Quantitative histomorphometric analysis of fractal dimension (D), lacunarity (E), epidermal thickness (F), epidermal junction density (G), and relative cell density (H). All measurements are expressed as the ratio of wound tissue to adjacent intact skin. Bars represent mean ± SD. (*p < 0.05; n = 4 animals per group).

Quantitative analysis demonstrated greater collagen deposition in WDE-treated wounds, as reflected by increased MT-derived collagen intensity (WDE: 0.86 ± 0.06 vs. sham: 0.69 ± 0.06; p = 0.008; Figure 4C.1). Consistent with enhanced tissue formation, eosin intensity measured from H&E sections was also increased following WDE treatment (WDE: 1.32 ± 0.17 vs. sham: 0.86 ± 0.28; p = 0.038; Figure 4C.2).

Morphometric analysis further identified differences in tissue architecture following WDE treatment. Relative lacunarity was higher in the WDE group (WDE: 1.06 ± 0.10 vs. sham: 0.65 ± 0.27; p = 0.048; Figure 4E), whereas epidermal thickness was reduced (WDE: 1.76 ± 0.13 vs. sham: 2.27 ± 0.34; p = 0.050; Figure 4F). Epidermal junction density was also lower in WDE- treated wounds (WDE: 4.88 ± 1.45 vs. sham: 7.45 ± 1.07; p = 0.032; Figure 4G), consistent with restoration of a more organized epidermal architecture. Relative cell density was increased following WDE treatment (WDE: 1.71 ± 0.35 vs. sham: 0.99 ± 0.09; p = 0.022; Figure 4H), indicating enhanced cellularity during tissue regeneration.

Relative fractal dimension did not differ between groups (WDE: 1.01 ± 0.02 vs. sham: 1.05 ± 0.03; p = 0.074; Figure 4D), suggesting that overall tissue complexity was comparable despite marked differences in collagen deposition and tissue organization.

Collectively, these findings are consistent with WDE promoting tissue remodeling, characterized by increased collagen deposition, enhanced cellularity, and improved restoration of tissue architecture.

### WDE enhances cellular responses associated with tissue regeneration

Immunofluorescence analysis at day 14 demonstrated that WDE enhanced multiple cellular processes involved in wound repair and tissue regeneration (Figure 5). Representative immunostaining revealed increased expression of markers associated with myofibroblast activation, angiogenesis, and macrophage remodeling in WDE-treated wounds compared with sham (Figure 5A-E).

**Figure 5.**
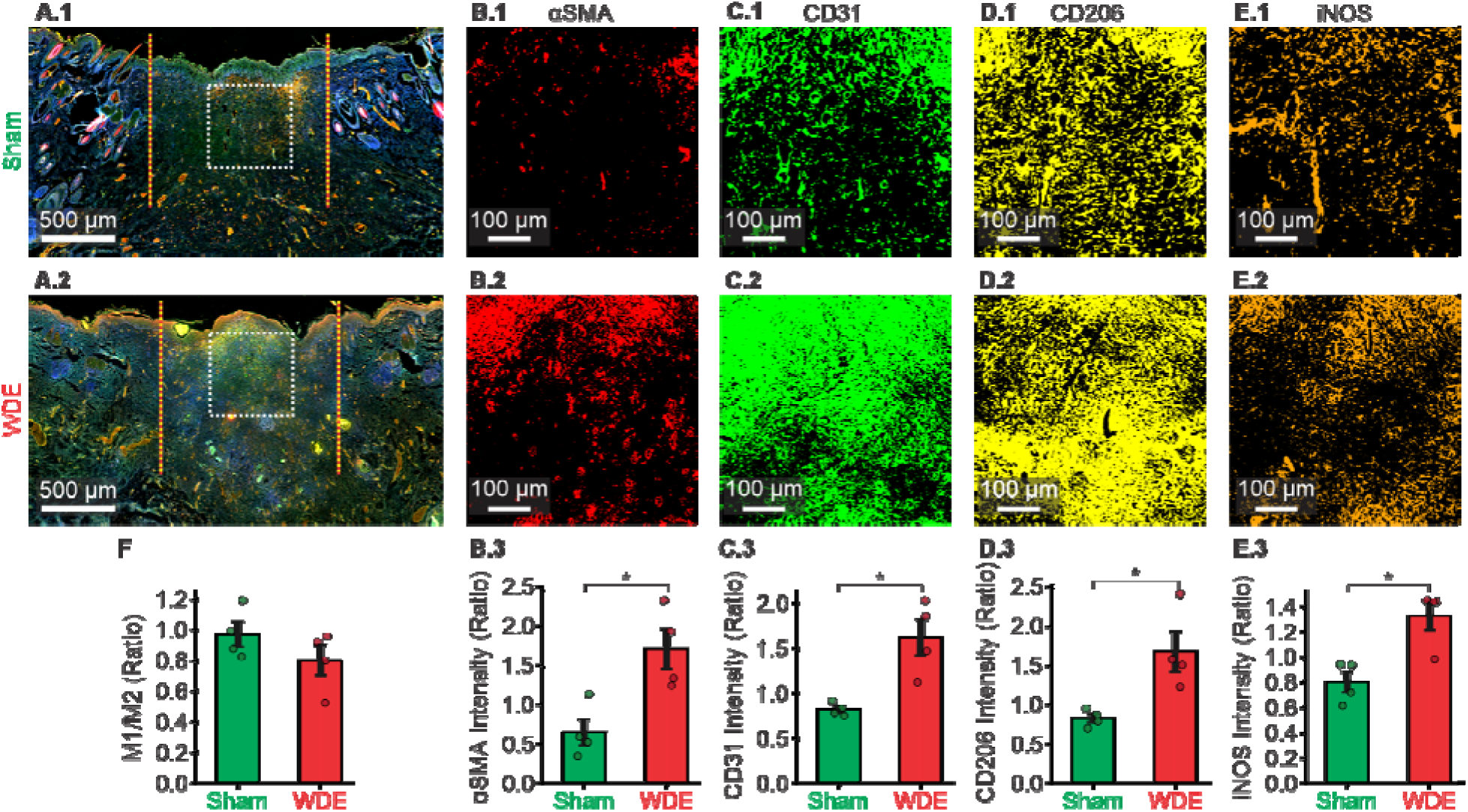
| Immunofluorescence characterization of wound repair after 14 days of electrotherapy. **(A.1-A.2)** Representative immunofluorescence images of sham- and WDE- treated wounds. Dashed orange lines indicate the approximate wound margins, and the white dashed squares denote regions selected for higher-magnification analysis. **(B-E)** Magnified images showing immunostaining for α-smooth muscle actin (αSMA; B.1-B.2), CD31 (C.1-C.2), CD206 (D.1-D.2), and inducible nitric oxide synthase (iNOS; E.1-E.2), with corresponding quantification of marker intensity (B.3-E.3) expressed as the ratio of wound tissue to adjacent intact skin. **(F)** Quantification of the macrophage polarization (M1/M2) index (iNOS/CD206), expressed as the ratio of wound tissue to adjacent intact skin. Bars represent mean ± SD (*p < 0.05; n = 4 animals per group).

Expression of α-smooth muscle actin (αSMA) was increased following WDE treatment (WDE: 1.71 ± 0.51 vs. sham: 0.65 ± 0.34; p = 0.017; Figure 5B), consistent with enhanced myofibroblast activation during wound remodeling. Likewise, CD31 expression was elevated (WDE: 1.62 ± 0.41 vs. sham: 0.82 ± 0.07; p = 0.027; Figure 5C), consistent with enhanced angiogenesis.

Macrophage-associated markers were enhanced by WDE treatment. Expression of the M2- associated marker CD206 was greater in the WDE group than in sham (WDE: 1.68 ± 0.51 vs. sham: 0.83 ± 0.11; p = 0.042; Figure 5D), while expression of the M1-associated marker inducible nitric oxide synthase (iNOS) was likewise elevated (WDE: 1.33 ± 0.23 vs. sham: 0.80 ± 0.16; p = 0.012; Figure 5E).

Despite the increase in both markers, the iNOS/CD206 ratio did not differ between groups (WDE: 0.80 ± 0.20 vs. sham: 0.97 ± 0.16; p = 0.233; Figure 5F), indicating increased macrophage marker expression without a detectable change in polarization balance.

Together, these findings demonstrate that WDE enhances coordinated cellular responses driving tissue repair, including increased expression of myofibroblast, angiogenesis, and macrophage markers.

### Computational modeling reveals localized electrical stimulation within the wound microenvironment

To investigate the electrical mechanisms underlying WDE-mediated wound healing, we developed the first cellular-scale computational model of wound electrotherapy incorporating representative skin microarchitecture (Figure 6). Electrical conductivities were assigned to resolved tissue structures, including the epidermis, dermis, vasculature, skin appendages, adipose tissue, and representative cellular components (Methods, Figure 6A). This heterogeneous conductivity distribution (10^-6^-1 S/m) was the substrate for cellular-scale predictions on stimulating current flow across the wound microenvironment.

**Figure 6.**
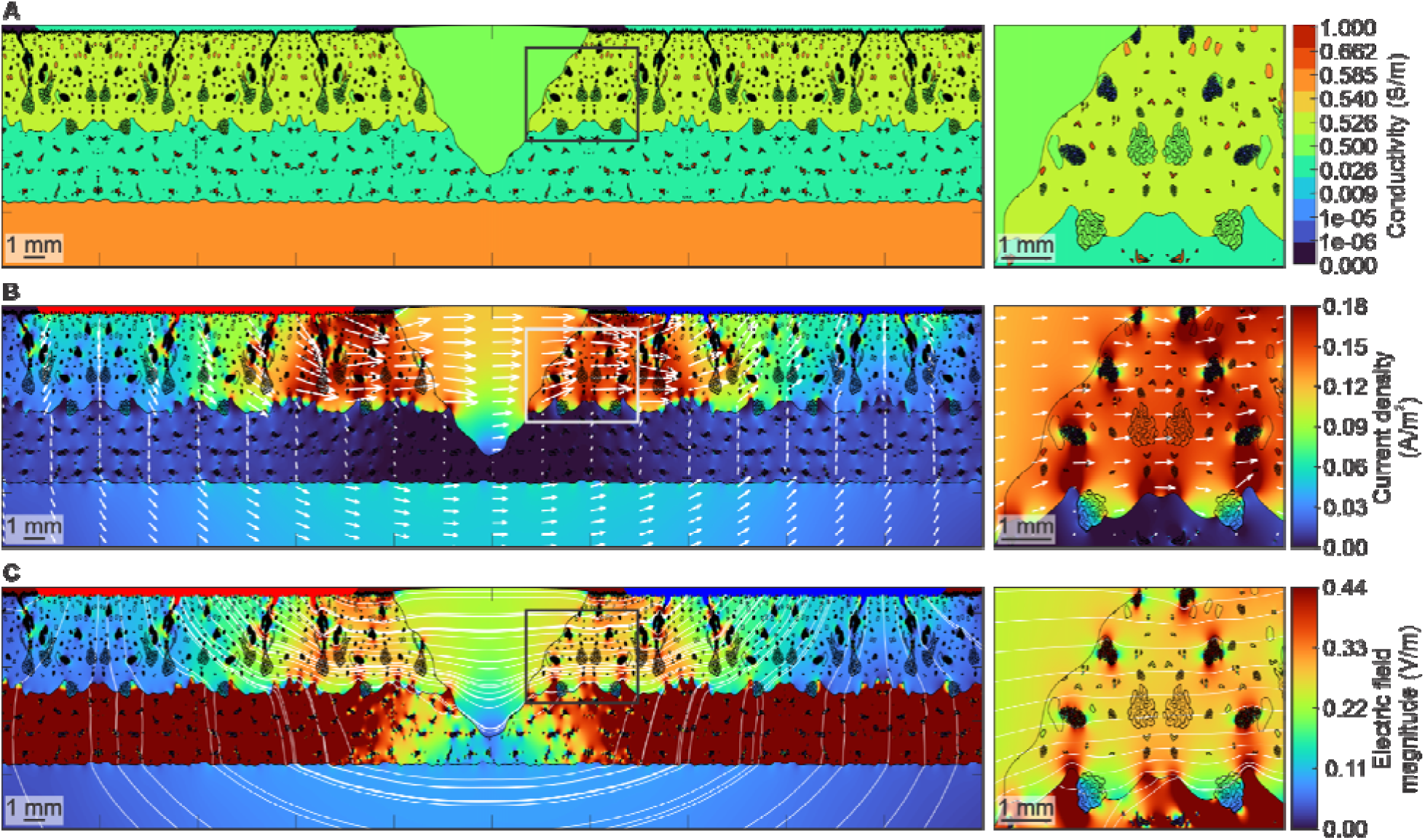
| Cellular-resolution computational model of electrical stimulation within the wound microenvironment. **(A)** Two-dimensional computational model of the microscopic tissue and cellular architecture surrounding the wound. Peri-wound stimulation is delivered through electrodes positioned on intact skin, avoiding direct contact with the wound bed. **(B)** Simulated current density distribution (linear scale) during peri-wound electrical stimulation. White arrows indicate current flow direction, with current preferentially concentrated and redirected tangentially along the wound margins. **(C)** Simulated electric field magnitude during stimulation. Local electric field concentrations occur around heterogeneous cellular and tissue structures. White streamlines indicate electric field direction. Right panels show magnified views of the highlighted regions in (A-C), illustrating localized current pathways and electric field distributions arising from microscale tissue heterogeneity.

Simulations predicted highly non-uniform current density throughout the tissue, with preferential current flow through electrically conductive structures and concentration (up to 0.18 A/m^2^) at tissue interfaces adjacent to the wound margins (Figure 6B). Corresponding electric field distributions were similarly shaped by both tissue- and cellular-scale heterogeneity (Figure 6C). Peak electric field magnitudes reached ∼0.30 V/m near the wound edge and exceeded 0.44 V/m at conductive skin appendages and tissue interfaces, including hair follicles, sweat glands, blood vessels, and adjacent cellular structures (Figure 6A-C, right panels). At the peri-wound region, both current density and electric field were tangential to the wound surface, analogous to endogenous injury currents [3,4].

These simulations - the first to resolve cellular-scale wound microarchitecture by leveraging BEM-FMM solvers - demonstrate that tissue electrical heterogeneity generates localized current pathways and electric field hotspots concentrated at wound boundaries and adjacent cellular structures. These findings provide a mechanistic framework linking externally applied electrical stimulation to the localized biophysical cues that drive tissue remodeling and regeneration.

## Discussion

This study demonstrates that a self-powered wearable disposable electrotherapy (WDE) platform delivers electrical stimulation comparable to a conventional CC system while significantly accelerating wound healing in a preclinical full-thickness skin wound model. WDE accelerated wound closure, enhanced tissue remodeling through increased collagen deposition and improved epidermal organization, and elicited regenerative cellular responses consistent with enhanced angiogenesis, myofibroblast activation, and macrophage modulation.

Although electrical stimulation has long been recognized to promote wound healing [7,10,31,32], clinical adoption remains limited by the need for wiring to external stimulators or self-contained devices with limited dose control [10,11]. Existing approaches disrupt the wound-bed or standards of care [10]. WDE addresses these barriers with specialized printed electrochemistry providing power and stimulation control [11] in mm-thick flexible patches that appear and are used like a conventional bandage.

Beyond accelerating wound closure, WDE enhanced tissue remodeling, increasing collagen deposition, cellularity, and epidermal organization. These findings are consistent with the coordinated roles of extracellular matrix deposition and cellular remodeling during the proliferative and remodeling phases of wound healing [1,6,33]. These structural changes were accompanied by increased αSMA and CD31 expression, indicating enhanced myofibroblast activity and angiogenesis [1], together with increased expression of both M1- and M2- associated macrophage markers consistent with greater macrophage recruitment [12]. Computational modeling further showed that tissue heterogeneity produces localized current pathways and electric field distributions in periwound tissue and appendages, providing a biophysical framework linking electrical stimulation to tissue repair. Collectively, these findings suggest WDE enhances multiple coordinated pathways involved in tissue regeneration [1,4,7].

WDE simplifies electrotherapy by integrating the power source, electrochemical dose control, stimulation electrodes, and wound dressing into a single disposable patch that eliminates external stimulators, connecting wires, electronics, and direct electrode contact with the wound bed while delivering therapeutic electrical stimulation by simple application across the wound. The modular device design (Figure 1C) facilitates therapy across wounds of varying size and anatomical location, enabling treatment across home and medically involved (e.g., post-surgical) settings. Clinical trials are warranted to test the application of WDE spanning accelerated wound healing, long-term scar quality, functional recovery, pain management, or chronic or diabetic wounds.

## Acknowledgments

The authors thank Mohammad Ahmad Sabri and Akon Undieh for their assistance with blinded manual wound boundary delineation and scoring, and Niranjan Khadka for his guidance and technical assistance with COMSOL-based computational modeling of electrical stimulation in the rodent wound model.

## Author contributions

Conceptualization: M.B., M.B.K.

Methodology: M.B.K., M.F.R., D.A.D., G.N.P., S.N.M.

Investigation: M.B.K., A.S.K., R.B., B.M., M.P.P.

Formal analysis: M.B.K., K.D.

Computational modeling: M.B.K., D.A.D., G.N.P., S.N.M.

Visualization: M.B.K., A.S.K., D.A.D., G.N.P., S.N.M., M.R.D.U.

Resources: M.F.R., M.B.

Supervision: M.B.

Project cc: M.B.K., M.B.

Writing—original draft: M.B.K., M.B. Writing—review & editing: All authors.

## References

1. H. N. Wilkinson, M. J. Hardman, Wound healing: cellular mechanisms and pathological outcomes. Open Biol 10, 200223 (2020).

2. G. C. Gurtner, S. Werner, Y. Barrandon, M. T. Longaker, Wound repair and regeneration. Nature 453, 314–321 (2008).

3. I. S. Foulds, A. T. Barker, Human skin battery potentials and their possible role in wound healing. Br. J. Dermatol. 109, 515–522 (1983).

4. M. Zhao, J. Penninger, R. R. Isseroff, Electrical Activation of Wound-Healing Pathways. Adv Skin Wound Care 1, 567–573 (2010).

5. R. Luo, J. Dai, J. Zhang, Z. Li, Accelerated Skin Wound Healing by Electrical Stimulation. Adv Healthc Mater 10, e2100557 (2021).

6. M. Zhao, H. Bai, E. Wang, J. V. Forrester, C. D. McCaig, Electrical stimulation directly induces pre-angiogenic responses in vascular endothelial cells by signaling through VEGF receptors. J. Cell Sci. 117, 397–405 (2004).

7. M. Rabbani, E. Rahman, M. B. Powner, I. F. Triantis, Making sense of electrical stimulation: A meta-analysis for wound healing. Ann. Biomed. Eng. 52, 153–177 (2024).

8. C. Wang, X. Jiang, H.-J. Kim, S. Zhang, X. Zhou, Y. Chen, H. Ling, Y. Xue, Z. Chen, M. Qu, L. Ren, J. Zhu, A. Libanori, Y. Zhu, H. Kang, S. Ahadian, M. R. Dokmeci, P. Servati, X. He, Z. Gu, W. Sun, A. Khademhosseini, Flexible patch with printable and antibacterial conductive hydrogel electrodes for accelerated wound healing. Biomaterials 285, 121479 (2022).

9. Y. Jiang, A. A. Trotsyuk, S. Niu, D. Henn, K. Chen, C.-C. Shih, M. R. Larson, A. M. Mermin-Bunnell, S. Mittal, J.-C. Lai, A. Saberi, E. Beard, S. Jing, D. Zhong, S. R. Steele, K. Sun, T. Jain, E. Zhao, C. R. Neimeth, W. G. Viana, J. Tang, D. Sivaraj, J. Padmanabhan, M. Rodrigues, D. P. Perrault, A. Chattopadhyay, Z. N. Maan, M. C. Leeolou, C. A. Bonham, S. H. Kwon, H. C. Kussie, K. S. Fischer, G. Gurusankar, K. Liang, K. Zhang, R. Nag, M. P. Snyder, M. Januszyk, G. C. Gurtner, Z. Bao, Wireless, closed-loop, smart bandage with integrated sensors and stimulators for advanced wound care and accelerated healing. Nat. Biotechnol. 41, 652–662 (2023).

10. Y. J. Cheah, M. R. Buyong, M. H. Mohd Yunus, Wound Healing with Electrical Stimulation Technologies: A Review. Polymers (Basel) 13, 3790 (2021).

11. M. FallahRad, K. Donnery, M. Belali Koochesfahani, Z. Chaudhry, R. Bhuiyan, B. Babaev, M. Saw, T. Liu, M. R. Diaz Uraga, M. Zaman, K. Saha, O. Velarde, A. Rddad, N. Khadka, M. Thahsin, A. Couzis, M. Bikson, Wearable disposable electrotherapy. Nat. Commun. 16, 9060 (2025).

12. R. Kaveti, M. A. Jakus, H. Chen, B. Jain, D. G. Kennedy, E. A. Caso, N. Mishra, N. Sharma, B. E. Uzunoğlu, W. B. Han, T.-M. Jang, S.-W. Hwang, G. Theocharidis, B. J. Sumpio, A. Veves, S. K. Sia, A. J. Bandodkar, Water-powered, electronics-free dressings that electrically stimulate wounds for rapid wound closure. Sci. Adv. 10, eado7538 (2024).

13. X. Sun, Y. Yang, Q. Liu, D. Zheng, C. Shao, Y. Wang, J. Lv, T. Yang, Y. Lu, Q. Ren, N. Chen, Accelerating wound healing with flexible zinc ion microbatteries coupled with endogenous electrical fields. Nano Energy 123, 109425 (2024).

14. G. Thakral, J. Lafontaine, B. Najafi, T. K. Talal, P. Kim, L. A. Lavery, Electrical stimulation to accelerate wound healing. Diabet Foot Ankle 4 (2013).

15. C. Khouri, S. Kotzki, M. Roustit, S. Blaise, F. Gueyffier, J.-L. Cracowski, Hierarchical evaluation of electrical stimulation protocols for chronic wound healing: An effect size meta-analysis. Wound Repair Regen 25, 883–891 (2017).

16. H. A. Thawer, P. E. Houghton, Effects of electrical stimulation on the histological properties of wounds in diabetic mice. Wound Repair Regen 9, 107–115 (2001).

17. K. Goyal, D. A. Borkholder, S. W. Day, Dependence of Skin-Electrode Contact Impedance on Material and Skin Hydration. Sensors (Basel) 22, 8510 (2022).

18. A. Grada, J. Mervis, V. Falanga, Research techniques made simple: Animal models of wound healing. J. Invest. Dermatol. 138, 2095–2105.e1 (2018).

19. K. Young, H. Morrison, Quantifying Microglia Morphology from Photomicrographs of Immunohistochemistry Prepared Tissue Using ImageJ. J. Vis. Exp. 136, e57648 (2018).

20. K. Fancher, J. M. Gardner, S. C. Shalin, Elastophagocytosis and interstitial granulomatous infiltrate are more common in extragenital vs genital lichen sclerosus. J Cutan Pathol 47, 903–912 (2020).

21. M. S. Abate, L. R. Battle, A. N. Emerson, J. M. Gardner, S. C. Shalin, Dermatologic Urgencies and Emergencies: What Every Pathologist Should Know. Arch Pathol Lab Med 143, 919–942 (2019).

22. E. H. Fulton, J. R. Kaley, J. M. Gardner, Skin Adnexal Tumors in Plain Language: A Practical Approach for the General Surgical Pathologist. Arch Pathol Lab Med 143, 832– 851 (2019).

23. M. Johnson, D. Walker, W. Galloway, J. M. Gardner, S. C. Shalin, Interface dermatitis along Blaschko’s lines. J Cutan Pathol 41, 950–954 (2014).

24. J. Hardin, J. M. Gardner, M. I. Colomé, P. Chévez-Barrios, Verrucous cyst with melanocytic and sebaceous differentiation: a case report and review of the literature. Arch Pathol Lab Med 137, 576–579 (2013).

25. R. Deivasigamani, N. N. M. Maidin, M. F. M. R. Wee, M. A. Mohamed, M. R. Buyong, Dielectrophoresis Prototypic Polystyrene Particle Synchronization toward Alive Keratinocyte Cells for Rapid Chronic Wound Healing. Sensors (Basel) 21, 3007 (2021)..

26. B. Tsai, H. Xue, E. Birgersson, S. Ollmar, U. Birgersson, Dielectrical Properties of Living Epidermis and Dermis in the Frequency Range from 1 kHz to 1 MHz. J Electr Bioimpedance 10, 14–23 (2019).

27. T. N. G. Adams, P. A. Turner, A. V. Janorkar, F. Zhao, A. R. Minerick, Characterizing the dielectric properties of human mesenchymal stem cells and the effects of charged elastin-like polypeptide copolymer treatment. Biomicrofluidics 8, 054109 (2014).

28. T. J. Faes, H. A. van der Meij, J. C. de Munck, R. M. Heethaar, The electric resistivity of human tissues (100 Hz-10 MHz): a meta-analysis of review studies. Physiol Meas 20, R1– 10 (1999).

29. L. Ortega, A. Llorella, J. P. Esquivel, N. Sabaté, Self-powered smart patch for sweat conductivity monitoring. Microsyst Nanoeng 5, 3 (2019).

30. J. H. Lee, Y. C. Yoon, H. S. Kim, J. Lee, E. Kim, C. Findeklee, U. Katscher, In vivo electrical conductivity measurement of muscle, cartilage, and peripheral nerve around knee joint using MR-electrical properties tomography. Sci. Rep. 12, 73 (2022).

31. S. N. Makarov, G. M. Noetscher, T. Raij, A. Nummenmaa, A Quasi-Static Boundary Element Approach With Fast Multipole Acceleration for High-Resolution Bioelectromagnetic Models. IEEE Trans Biomed Eng 65, 2675–2683 (2018).

32. D. A. Drumm, G. M. Noetscher, H. Oppermann, J. Haueisen, Z.-D. Deng, S. N. Makaroff, Charge Based Boundary Element Method with Residual Driven Adaptive Mesh Refinement for High Resolution Electrical Stimulation Modeling, bioRxiv (2026). 10.64898/2026.03.11.711201.

33. S. N. Makarov, L. Golestanirad, W. A. Wartman, B. T. Nguyen, G. M. Noetscher, J. P. Ahveninen, K. Fujimoto, K. Weise, A. R. Nummenmaa, Boundary element fast multipole method for modeling electrical brain stimulation with voltage and current electrodes. J Neural Eng 18 (2021).

