## Supplementary Figure 1 for "Self-powered electronics-free Wearable Disposable Electrotherapy (WDE) platform for accelerated wound healing"

**bioRxiv**

**Mojtaba Belali Koochesfahani et al*.***

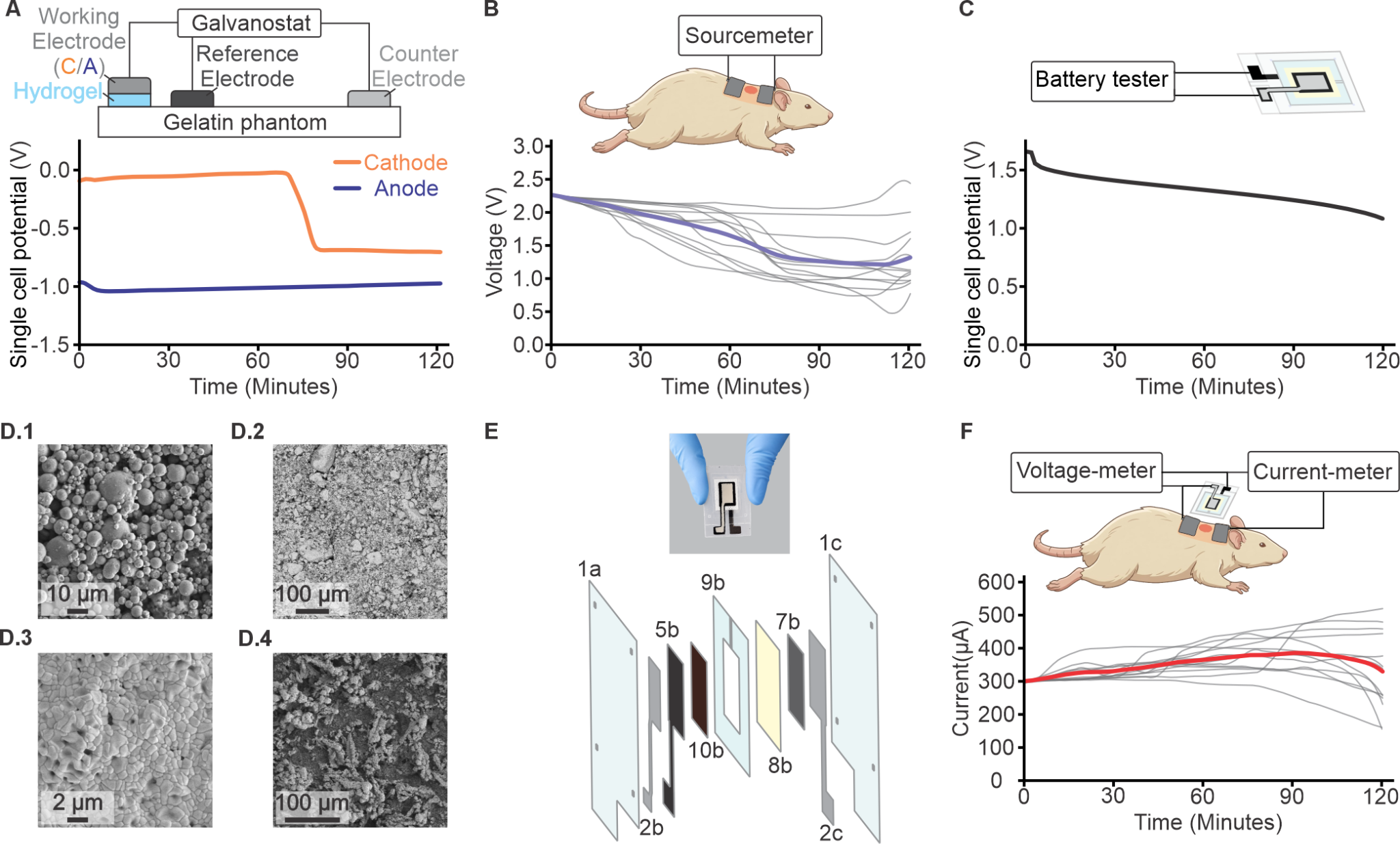

**Supplementary Figure 1 | Wearable Disposable Electrotherapy device design for rat model. (A)** Electrochemical characterization of stimulation electrodes by galvanostatic discharge (300 µA) using saline-soaked foam. **(B)** Experimental setup for in vivo constant current (CC) stimulation (300 µA) and voltage recording during dose delivery (gray: individual tests; purple: group’s mean; n = 12). **(C)** Experimental setup and galvanostatic discharge profile of the printed battery cell (300 µA, 120 minutes). Discharge profile of the printed battery cell. **(D)** Scanning electron microscopy (SEM) of the printed battery anode (D.1), printed battery cathode (D.2), Ag/AgCl stimulation electrode (D.3), and zinc stimulation electrode (D.4), illustrating the surface microstructure of the printed electrochemical components. **(E)** Photograph of the fabricated printed electrochemical cell with an exploded view of the multilayer architecture. **(F)** Experimental setup for current and voltage recording during dose application using a WDE patch (gray: recorded current of sessions; red: group’s mean; n = 13).
